# DNA methylation subtypes inferred from transcriptomes guide sunitinib and immunotherapy selection in renal cancer

**DOI:** 10.1101/2024.11.05.622095

**Authors:** Florian Jeanneret, Sarah Schoch, Pedro Ballester, Stefan N Symeonides, Alexander Laird, Håkan Axelson, Delphine Pflieger, Christophe Battail

## Abstract

Responses to targeted and immune checkpoint therapies in metastatic clear cell renal cell carcinoma (ccRCC) are heterogeneous, and no predictive biomarker is available. We performed a meta-analysis of DNA methylation profiles from 727 ccRCC tumors, benchmarked four published signatures, and identified a hyper-methylated subtype (15% of tumors) associated with shorter overall survival. Integration of 589 matched transcriptomes showed that hyper-methylated tumors were more proliferative, depleted in endothelial tip cells, and overexpressed DNMT3B while underexpressing TET2. We then developed a gene expression classifier based on nine gene ratios that predicted methylation subtypes in an independent cohort (AUC-ROC = 0.96). In four treated cohorts, predicted hyper-methylated tumors had shorter progression-free survival on sunitinib but not on immune checkpoint-based regimens. Hyper-methylated primary tumors responded better to atezolizumab plus bevacizumab than to sunitinib, whereas hypo-methylated metastases benefited more from sunitinib. Methylation subtypes may thus help guide treatment selection in metastatic ccRCC.

## Introduction

Renal cell carcinoma (RCC) accounts for the large majority of kidney cancers, which represented approximately 435,000 new cases and 156,000 deaths worldwide in 2022 ^1^. Clear cell RCC (ccRCC) is the most common histological subtype, accounting for approximately 80% of cases, whereas papillary and chromophobe RCC represent most of the remainder ^2^. Partial or radical nephrectomy remains the only curative treatment for localized disease ^2,3^ and adjuvant pembrolizumab has improved outcomes in patients at high risk of recurrence ^4^. In the metastatic setting, tyrosine kinase inhibitors (TKI) targeting the VEGF pathway, mTOR inhibitors, and more recently immune checkpoint inhibitors (ICI) alone or combined with TKI, have substantially prolonged survival ^5,6^. Yet responses remain highly variable: many patients do not achieve durable benefit while being exposed to serious treatment-related toxicities ^7–12^. No predictive molecular biomarker is currently approved to guide treatment selection in metastatic ccRCC, which remains a major unmet clinical need.

Large-scale genomic, transcriptomic, proteomic and epigenomic profiling has greatly advanced our understanding of ccRCC biology ^13–17^, but the determinants of treatment outcome remain poorly understood. Biomarkers established in other tumor types have shown limited value in this disease. PD-L1 expression assessed by immunohistochemistry is an FDA-approved companion diagnostic for ICIs in several cancers ^18^, yet it predicts response imperfectly ^19^ and is not used to select therapy in RCC. Likewise, tumor mutational burden (TMB) correlates with ICI response in non-small cell lung cancer (NSCLC) and melanoma, but its predictive value is limited in ccRCC, which harbors a relatively low mutation rate ^20–22^. Transcriptomic signatures have also been proposed, including the JAVELIN Renal 101 signature ^22^ and the IMmotion151 molecular subsets ^10^ in ccRCC, and the MIAS^23^, GEP ^24^, MHC-I/II^25^ and IMPRES^26^ scores in melanoma. However, none has been adopted in routine practice ^27,28^, although the BIONIKK trial demonstrated the feasibility of subtype-driven treatment selection in ccRCC ^29^. Epigenomic approaches are also emerging: profiling of histone modifications in sarcomatoid RCC identified transcription factors, such as FOSL1, whose expression stratified patients by clinical outcome ^30^.

DNA methylation provides a complementary layer of information. Several studies identified two or three DNA methylation subtypes of ccRCC with distinct survival ^14,31,32^, and methylation-based prognostic scores have been proposed ^32–34^. However, their performance in independent cohorts has been inconsistent, and none was designed to predict response to targeted therapies or ICIs. Tumor DNA methylation has also been associated with ICI response in NSCLC, melanoma and ccRCC ^35–37^. Beuselinck et al. linked the ccrcc1 and ccrcc4 subgroups, which display a CpG island methylator phenotype, to poor sunitinib response ^38^. More recently, Lu et al. developed methylation- and expression-based classifiers in ccRCC patients treated with ICI or TKI ^37^, but the predicted subtypes were not associated with progression-free survival in independent cohorts.

A major obstacle to evaluating DNA methylation as a predictive biomarker is that tumor samples from clinical trials have mostly been profiled by transcriptomics ^8,22,24,26^, whereas methylation data are rarely available. Inferring methylation subtypes from gene expression data can overcome this limitation and offers a cost-effective strategy that is readily translatable to the clinic. Here, we performed a meta-analysis of DNA methylation profiles from 727 ccRCC tumors to investigate determinants of treatment response in metastatic disease. We first compared published methylation-based classification methods to select the most prognostic one. Using tumors profiled by both methylation arrays and transcriptomics, we then characterized differences between subtypes in tumor microenvironment composition, pathway activity and transcription factor activity. Next, we developed a simple gene-ratio classifier that predicts methylation subtypes from gene expression data. Finally, applying this classifier to four cohorts of treated patients, three of them from clinical trials, we show that methylation subtypes are associated with response to sunitinib and may inform the choice between TKI and ICI-based combinations.

## Results

### DNA methylation subtypes stratify patient survival across a ccRCC meta-cohort

Four previous studies classified ccRCC tumors from methylation array data. Two used unsupervised clustering (Arai et al.^31^, Sato et al.^14^) and two used supervised classification (Evelönn et al.^32^, Lu et al.^37^). Although these methods relied on different sets of CpG probes, all four associated a hyper-methylated profile with poor prognosis. To identify the most informative classification, we applied the four methods to a single unified cohort.

We collected methylation array data for 757 ccRCC samples from four data sources. After removal of duplicated patients and quality filtering, the harmonized and batch-corrected dataset comprised 727 samples, 589 of which had matched gene expression data from three sources (Fig. S1). Age, sex and clinical stage were similarly distributed across the four datasets (Table S1). Some signature probes were either absent from one of the Illumina 450K and EPIC arrays or had too many missing values. Only subsets of the original signatures could therefore be used: 531 of 801 probes for Arai et al., 1,006 of 1,672 for Sato et al. and 148 of 174 for Evelönn et al. (Fig. S1). Classifications obtained with these reduced probe sets were concordant with those obtained with the complete signatures (Tables S2-S4), indicating limited signal loss.

The Arai et al., Sato et al., Evelönn et al. and Lu et al. methods classified 267, 109, 274 and 242 of the 727 tumors as hyper-methylated, respectively (Fig. 1a, Table S5). In all four methods, the hyper-methylated group was associated with shorter OS (two-sided log-rank P = 1.9 × 10⁻⁴, 1.6 × 10⁻¹⁰, 1.0 × 10⁻⁵ and 2.2 × 10⁻⁹ for the Arai et al., Sato et al., Evelönn et al. and Lu et al. methods, respectively; Fig. 1B-E). Although discrimination was comparable across methods (Harrell’s C-index 0.62-0.67; Table S16), the Sato et al. method identified the smallest hyper-methylated group (109 of 727 tumors, 15%) with the highest risk of death (HR 4.61, 95% CI 2.76-7.73). We therefore retained it as the reference classification for all subsequent analyses.

**Figure 1:**
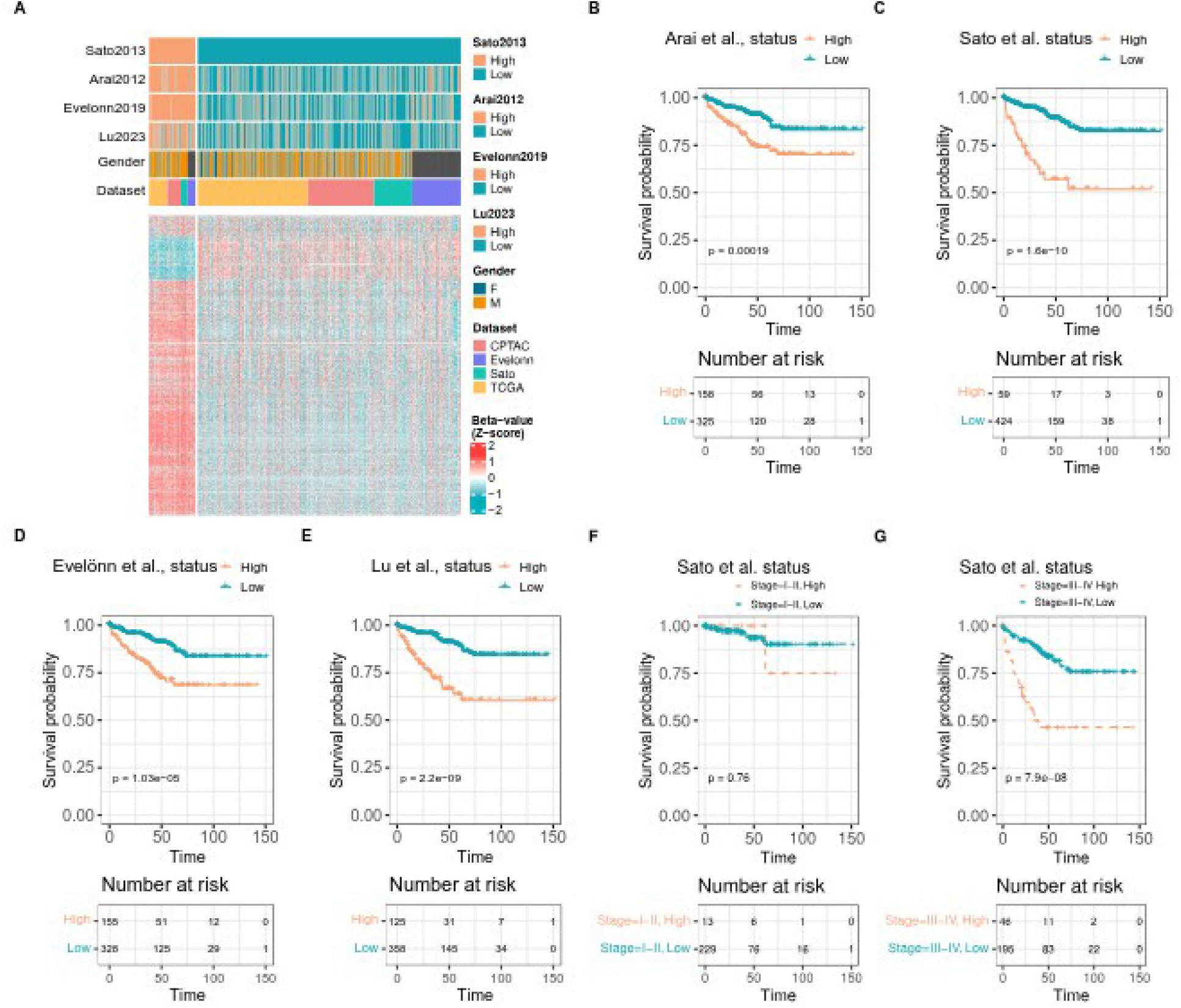
Association of ccRCC DNA methylation subtypes with overall survival. **A.** Hierarchical clustering of 727 ccRCC samples from the unified cohort (TCGA, CPTAC, Sato et al. and Evelönn et al. datasets) into hyper-methylated (High) and hypo-methylated (Low) subtypes according to the Sato et al. method. The heatmap shows the beta values of the 1,006 CpG probes of the Sato et al. signature retained after data harmonization. Kaplan-Meier curves of overall survival (OS) between methylation subtypes defined by **B.** the Arai et al., **C.** the Sato et al., **D.** the Evelönn et al. and **E.** the Lu et al. methods. **F.** OS analysis between Sato et al. subtypes in patients tumors with **F.** stages I and II and **G.** stages III and IV samples. OS is expressed in months. P values were calculated with two-sided log-rank tests.

Both methylation subtypes showed higher methylation levels than matched normal adjacent tissues in the TCGA dataset (450k Illumina array) (Fig. S2). The terms "hyper-methylated" and "hypo-methylated" therefore describe relative methylation levels among tumors, not relative to normal kidney.

Methylation subtypes were not associated with OS in patients with stage I-II tumors (Fig. 1F). In patients with stage III-IV disease, however, hyper-methylated tumors were associated with significantly shorter OS (Fig. 1G). In a multivariable Cox model, tumor stage and methylation subtype were both independently associated with OS (P < 1e-3, Fig. S3). Although the sex ratio was similar across cohorts, the hyper-methylated subtype included fewer tumors from women than the hypo-methylated subtype (16% versus 28%; Pearson’s chi-squared, P = 0.0035; Table S6). Finally, partitioning tumors into three methylation clusters instead of two did not improve prognostic stratification (Fig. S4).

### Hypermethylated tumors are proliferative, depleted in endothelial tip cells and show deregulation of DNMT3B and TET2

To understand the biological basis of the poor prognosis associated with hyper-methylation, we characterized the 589 tumors from the TCGA, CPTAC and Sato et al. datasets with matched methylation and gene expression data at the level of cellular composition, pathway activity, transcriptional regulation and DNA methylation machinery.

Because the composition of the tumor microenvironment (TME) influences both prognosis and treatment response ^39–41^, we first compared cell type fractions between methylation subtypes. We estimated the fractions of 21 cell types by CIBERSORTx deconvolution, using a ccRCC single-cell reference refined in our previous work ^42^ (Tables S7-S9). Compared with hypo-methylated tumors, hyper-methylated tumors were enriched in cycling tumor cells. They were depleted in tumor cells, CA9-high tumor cells, stromal cells and endothelial tip cells (Fig. 2A, B). In contrast, none of the 15 immune cell fractions differed between subtypes (Table S10). A transcriptomic score of tertiary lymphoid structures (TLS) ^39^ was also similar in both groups (two-sided Wilcoxon rank-sum test, P > 0.05; Fig. S5, Table S11). The two subtypes thus displayed comparable immune compositions but distinct tumor cell states and endothelial content.

**Figure 2:**
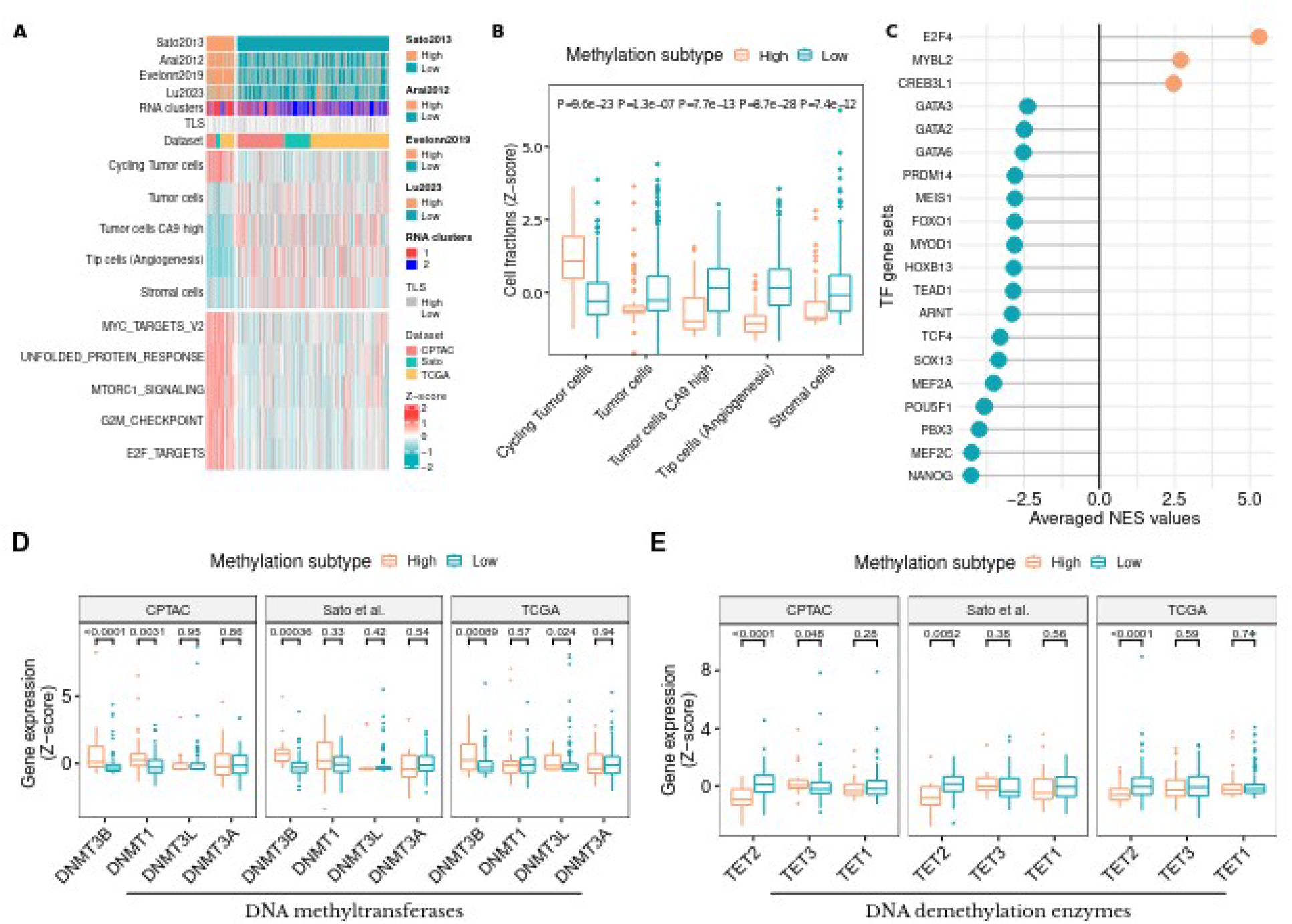
Transcriptomic characterization of ccRCC DNA methylation subtypes. Analyses were performed separately in the TCGA, CPTAC and Sato et al. datasets (589 tumors with matched methylation and gene expression data), using the Sato et al. subtypes (High, hypermethylated; Low, hypomethylated). **A.** Heatmap of cell type fractions estimated by CIBERSORTx deconvolution and of ssGSEA scores of MSigDB hallmark gene sets. Tumors are ordered by methylation subtype. "RNA clusters" indicates groups obtained by consensus clustering of gene expression data. **B.** Cell type fractions differing significantly between methylation subtypes (Wilcoxon rank-sum tests). **C.** Transcription factor (TF) activities inferred with the VIPER algorithm from DoRothEA regulons. Positive and negative normalized enrichment scores (NES) indicate higher and lower activity in hype-rmethylated tumors, respectively. The 20 most significant TFs are shown. **D.** Expression of DNA methyltransferases between subtypes **E.** Expression of DNA demethylation enzymes between subtypes. In **B**, **D** and **E**, P values were calculated with two-sided Wilcoxon rank-sum tests and adjusted by the Benjamini-Hochberg method. In **B**, P values from the three datasets were combined by Stouffer’s method.

We next asked which biological programs distinguished the two subtypes, using single-sample gene set enrichment analysis (ssGSEA) of the 50 hallmark and 189 oncogenic signature gene sets from MsigDB ^43,44^. Sixteen hallmark gene sets differed significantly between subtypes in all three datasets (Fig. 2A, Table S12). The five most significant were all up-regulated in hyper-methylated tumors, with Stouffer-combined adjusted P values as follows: unfolded protein response, P < 1e-21; mTORC1 signaling, P < 1e-21; MYC targets, P < 1e-22; G2/M DNA damage checkpoint, P < 1e-21; E2F targets, P < 1e-21. Among oncogenic signatures, 30 gene sets were significant across the three datasets (Fig. S6, Table S13). Genes upregulated by E2F1 were enriched in hyper-methylated tumors (corrected Stouffer P < 1e-18). Genes upregulated by GLI1, an effector of Hedgehog signaling, were depleted (GLI1_UP.V1_UP, corrected Stouffer P < 1e-19). Two independent mTOR perturbation signatures supported increased mTOR activity in hypermethylated tumors: genes expressed at higher levels in placebo-than in everolimus-treated mouse prostate (MTOR_UP.V1_UP) were enriched, whereas genes induced by rapamycin in CEM-C1 T-lymphoblastic cells (MTOR_UP.N4.V1_DN) were depleted (both corrected Stouffer P < 1e-17; Fig. S6). These results are consistent with the activation of the mTORC1 hallmark gene set.

### Hyper-methylated tumors show reduced activity of homeobox and GATA transcription factors

To identify transcriptional regulators that might drive these differences, we inferred transcription factor (TF) activities from the expression of their target genes with the VIPER algorithm ^45^. We used DoRothEA regulons ^46^ with confidence levels A-C and at least 15 measured targets. P values were combined across the three cohorts by Stouffer’s method and normalized enrichment scores (NES) by averaging. Twenty-two TFs differed significantly between subtypes: 3 showed higher and 19 showed lower activity in hyper-methylated tumors (Fig. 2C, Table S14). Consistent with the pathway analysis, the cell cycle regulator E2F4 was among the most activated TFs in hyper-methylated tumors (corrected Stouffer P < 1e-16). Conversely, the homeobox TFs NANOG, PBX3 and POU5F1 were among the most repressed (corrected Stouffer P < 1e-10, P < 1e-10 and P < 1e-8, respectively). These TFs have been implicated in stemness and tumorigenesis ^47–49^. The reduced activity of NANOG was not mirrored by lower mRNA levels in the TCGA (two-sided Wilcoxon rank-sum test, P=0.45) and CPTAC (two-sided Wilcoxon rank-sum test, P = 0.12) datasets, although its expression differed in the Sato et al. dataset (two-sided Wilcoxon rank-sum test, P = 6.9e-3; Fig. S7). CREB3L1, an endoplasmic reticulum stress-associated TF, showed higher activity in hyper-methylated tumors, in line with the activation of the unfolded protein response ^50^ (Fig. 2A, C). Finally, GATA2, GATA3 and GATA6 showed lower activity in hyper-methylated tumors. Reduced expression of GATA factors has been associated with tumor aggressiveness and poor outcome in ccRCC ^51–54^.

Finally, because DNA methylation levels result from the balance between DNA methyltransferase (DNMT) and demethylase activities, we examined whether the expression of these enzymes differed between subtypes. Among DNMTs, only DNMT3B, which mediates de novo methylation, was over-expressed in hyper-methylated tumors in all three datasets (two-sided Wilcoxon rank-sum tests, P < 0.001; Fig. 2D). DNMT1 (maintenance methylation), DNMT3A and the regulatory cofactor DNMT3L were not differentially expressed. Among the ten-eleven translocation (TET) demethylases, TET2 was consistently under-expressed in hyper-methylated tumors (two-sided Wilcoxon rank-sum tests, P < 0.01; Fig. 2E), whereas TET1 and TET3 were not. Increased DNMT3B and decreased TET2 expression may thus jointly contribute to the hyper-methylated phenotype. Overall, hyper-methylated ccRCCs combined four features: higher proliferative activity, a larger fraction of cycling tumor cells, reduced endothelial tip cell content, and deregulation of two key DNA methylation enzymes. These features may underlie their poorer survival.

### A gene-ratio classifier accurately predicts methylation subtypes from gene expression data

Hyper-methylated tumors were associated with shorter OS in advanced (stage III-IV) disease (Fig. 1G). We therefore asked whether methylation subtypes also influence response to systemic therapies. Because methylation profiling is rarely performed in clinical trials, we first developed a classifier that infers methylation subtypes from gene expression data.

Gene ratios are robust predictors of molecular subtypes across profiling platforms ^55–57^. We therefore used the Cross-Platform Omics Prediction (CPOP) algorithm ^58^ algorithm, which builds prediction models from gene ratios that are stable across training datasets. The model was trained on the TCGA (RNA-seq) and Sato et al. (microarray) datasets and evaluated on the independent CPTAC dataset. Input features were selected by grid search: 40 genes, drawn from the top 125 differentially expressed genes shared by both training datasets, were provided to CPOP, which retained nine stable gene ratios. Given the class imbalance, the model was optimized to minimize the false-positive rate, that is, the misclassification of hypo-methylated tumors as hyper-methylated.

In CPTAC, the classifier achieved an AUC-ROC of 0.96, a Matthews correlation coefficient (MCC) of 0.67 (95% CI 0.52-0.81), a specificity of 0.97 (0.94-0.99) and a sensitivity of 0.63 (0.45-0.77) (Table S15). Predicted hyper-methylated tumors were highly reliable (positive predictive value 0.83), whereas about one third of hyper-methylated tumors were classified as hypo-methylated. Like the subtypes defined directly from methylation array data (Fig. 3A), the predicted subtypes were associated with OS (two-sided log-rank P < 0.01; Fig. 3B).

**Figure 3:**
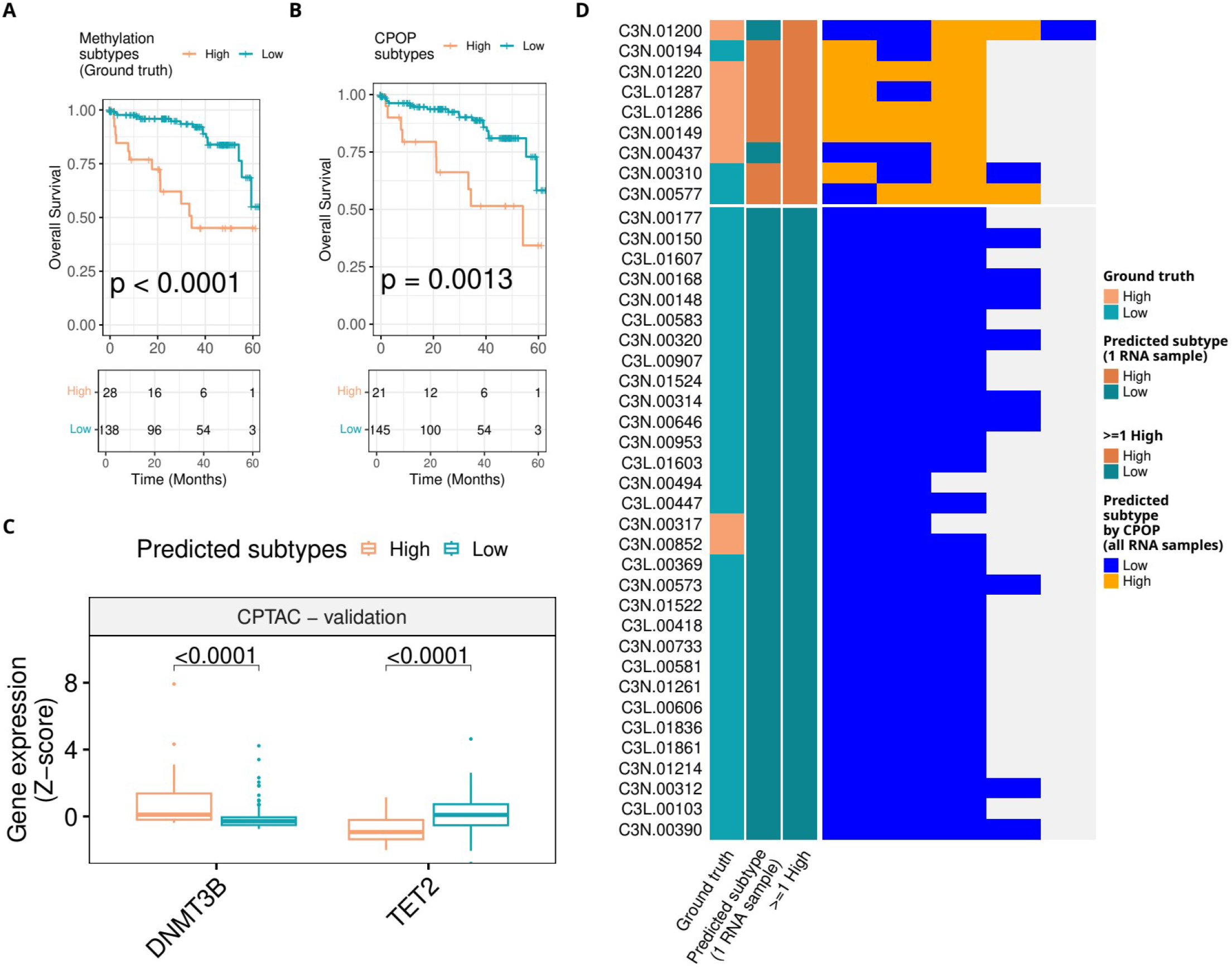
Prediction of DNA methylation subtypes from gene expression data with the CPOP classifier. Kaplan-Meier curves of OS according to methylation subtypes defined from methylation array data with **A.** the Sato et al. method (reference classification) or **B.** predicted with the CPOP classifier. P values were calculated with two-sided log-rank tests. **C.** Expression of DNMT3B and TET2 between predicted subtypes (two-sided Wilcoxon rank-sum tests). **D.** Impact of intratumoral heterogeneity (ITH) on predictions, assessed on 131 samples from multiple regions of the primary tumor of 40 CPTAC patients (at least two regions per patient). Patients are grouped according to whether at least one region was classified as hyper-methylated from methylation array data (≥1 High) or not.

We next assessed the impact of intratumoral heterogeneity (ITH) on predictions, using 131 samples from multiple regions of the primary tumor of 40 CPTAC patients (at least two regions per patient). Patients were grouped according to whether at least one region was classified as hyper-methylated from methylation array data (Fig. 3D). Predictions were largely concordant with this reference: most patients with at least one hyper-methylated region had two or more regions predicted as hyper-methylated, and only two had no region predicted as such. Several tumors nevertheless showed discordant predictions across regions, suggesting that sampling multiple regions could improve the robustness of patient classification.

### Predicted hyper-methylated tumors respond poorly to sunitinib but not to immune checkpoint-based therapies

We applied the classifier, without additional normalization, to four independent cohorts of patients with advanced ccRCC: the Beuselinck et al. cohort ^38^, treated with sunitinib (n = 51); the CheckMate cohort analyzed by Braun et al. ^11^, treated with nivolumab (n = 180); the IMmotion151 trial (Motzer et al. ^10^, n = 823); and the IMmotion150 trial (Banchereau et al. ^59^, n = 247). In IMmotion151 and IMmotion150, patients received sunitinib or atezolizumab plus bevacizumab (anti-PD-L1/anti-VEGF), and in IMmotion150 also atezolizumab alone. We analyzed primary tumors and metastases separately and compared progression-free survival (PFS) between predicted subtypes.

In patients treated with sunitinib, predicted hyper-methylated tumors were associated with significantly shorter PFS. This held both in primary tumors from the three sunitinib-treated cohorts (Fig. 4A-C) and in metastases from IMmotion151 (Fig. 4D). By contrast, predicted subtypes were not significantly associated with PFS in patients treated with nivolumab, atezolizumab or atezolizumab plus bevacizumab, whether in primary tumors or metastases (Fig. 4E-J).

**Figure 4:**
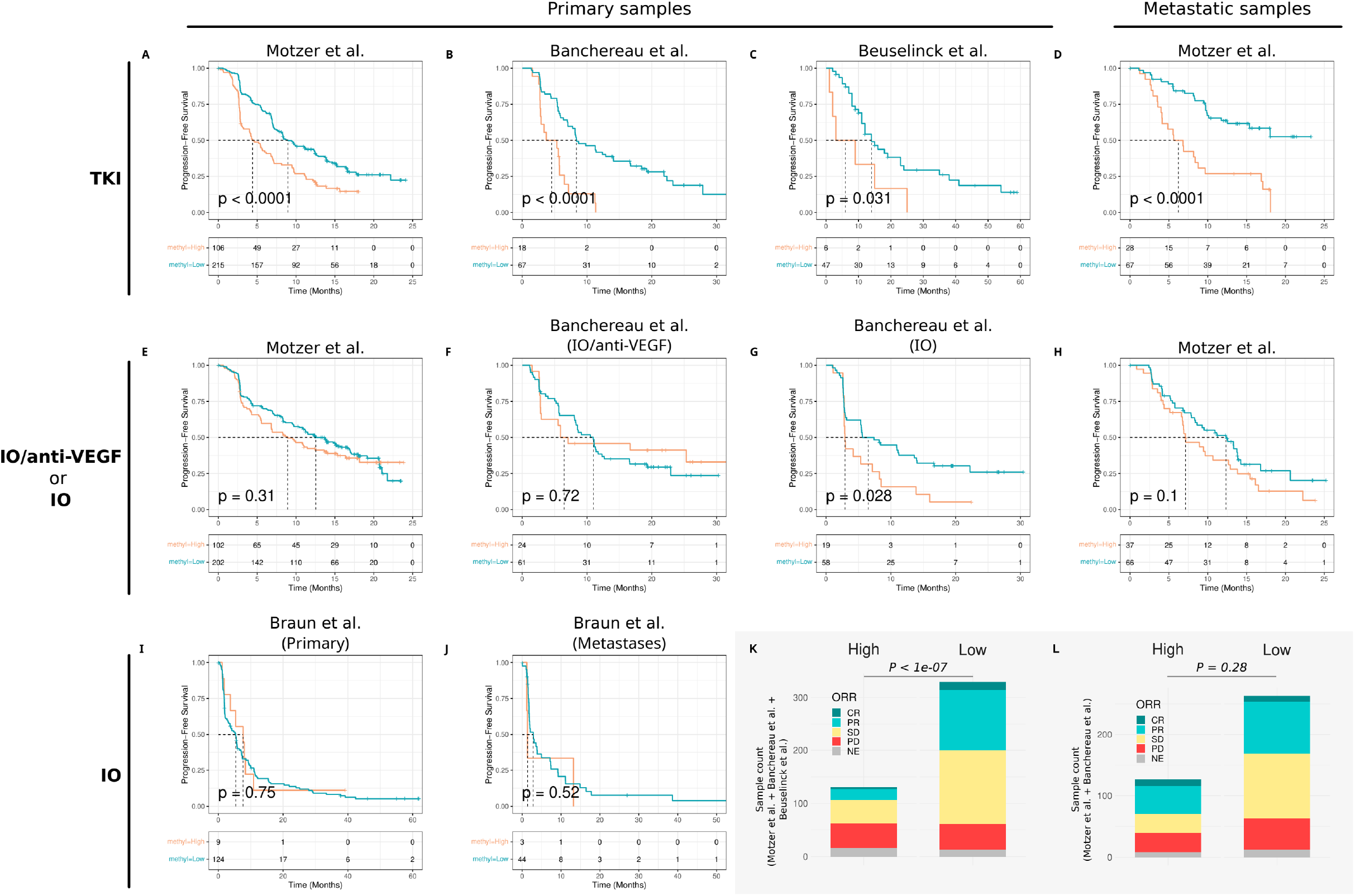
Association of predicted methylation subtypes with response to sunitinib and immune checkpoint-based therapies. Kaplan-Meier curves of progression-free survival (PFS) according to methylation subtypes predicted with the CPOP classifier. **A-D**, Sunitinib-treated patients: primary tumors from **A.** IMmotion151 (Motzer et al.), **B.** IMmotion150 (Banchereau et al.), and **C.** the Beuselinck et al. cohort; **D.** metastases from IMmotion151. **E-J**, Patients treated with immune checkpoint-based regimens: primary tumors treated with **E.** atezolizumab plus bevacizumab in IMmotion151 and **F.** IMmotion150; **G.** primary tumors treated with atezolizumab alone in IMmotion150; **H.** metastases treated with atezolizumab plus bevacizumab in IMmotion151; **I.** primary tumors and **J.** metastases treated with nivolumab in the CheckMate 009, 010 and 025 trials (Braun et al.). P values in **A-J** were calculated with two-sided log-rank tests. Distribution of RECIST response groups between predicted subtypes in pooled **K.** sunitinib-treated and **L.** atezolizumab plus bevacizumab-treated patients, restricted to complete response (CR), partial response (PR) and progressive disease (PD). P values were calculated with Pearson’s Chi-squared tests.

We then pooled all sunitinib-treated patients and all patients treated with atezolizumab plus bevacizumab. The distribution of best objective responses (RECIST) differed between predicted subtypes in the sunitinib group, with poorer outcomes among hyper-methylated tumors (Pearson’s Chi-quared test, P < 0.01; Fig. 4K). No such difference was observed in the atezolizumab plus bevacizumab group (Fig. 4L).

### Methylation subtypes may inform the choice between sunitinib and atezolizumab plus bevacizumab

IMmotion151 and IMmotion150 randomized patients between sunitinib and atezolizumab plus bevacizumab, and IMmotion150 also included an atezolizumab monotherapy arm ^60,61^. This design allowed us to compare treatments within each predicted subtype.

In primary tumors, the benefit of combination therapy depended on the methylation subtype. Among patients with hyper-methylated primary tumors, atezolizumab plus bevacizumab was associated with longer PFS than sunitinib in both trials, and also than atezolizumab alone in IMmotion150 (Fig. 5A, B). By contrast, PFS was similar across treatment arms among patients with hypo-methylated primary tumors (Fig. 5C, D). A different pattern emerged in metastatic samples: patients with hyper-methylated metastases had similar PFS with either treatment, whereas those with hypo-methylated metastases had significantly longer PFS with sunitinib (log rank P <0.01; Fig. 5E, F).

**Figure 5:**
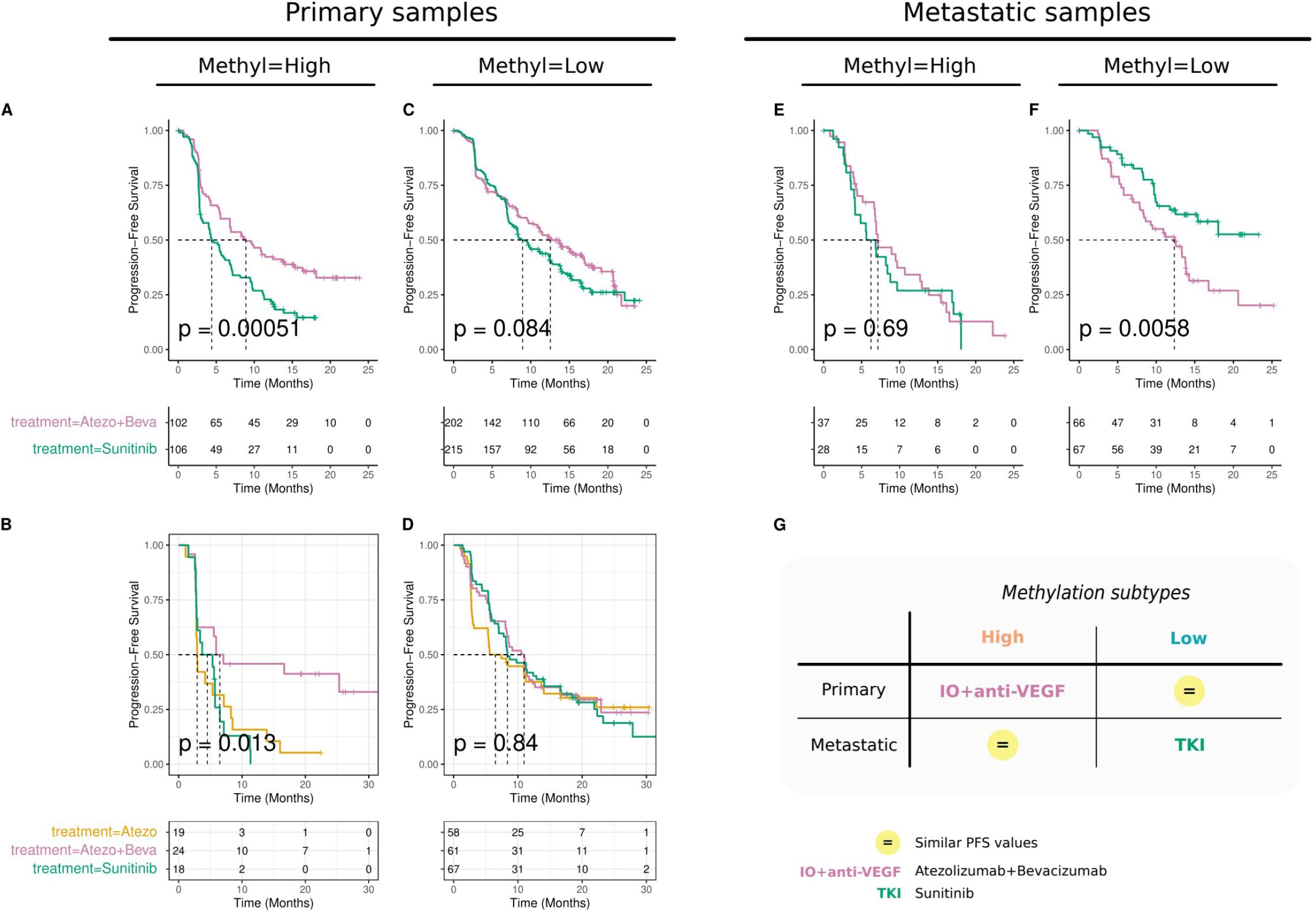
Comparison of sunitinib and atezolizumab-based therapies within predicted methylation subtypes. Kaplan-Meier curves of PFS by treatment arm in the randomized trials IMmotion150 and IMmotion151. Methylation subtypes were predicted with the CPOP classifier. **A.** Hyper-methylated (High) primary tumors in IMmotion151. **B.** Hyper-methylated primary tumors (High) in IMmotion150. **C.** Hypo-methylated (Low) primary tumors in IMmotion151. **D.** Hypo-methylated (Low) primary tumors in IMmotion150. **E.** Hyper-methylated metastases in IMmotion15. **F.** Hypo-methylated (f) metastases in IMmotion151. P values were calculated with two-sided log-rank tests. **G.** Putative framework for treatment selection according to predicted methylation subtype and sample type. Atezo, atezolizumab; Atezo+Beva, atezolizumab plus bevacizumab.

Methylation subtypes are thus associated with treatment response in advanced ccRCC, with differences between primary tumors and metastases. These results support a hypothesis-generating framework: atezolizumab plus bevacizumab would be preferred for patients with hyper-methylated primary tumors, and sunitinib for patients with hypo-methylated metastases (Fig. 5G). This framework will require prospective validation.

## Discussion

In this work, we assembled DNA methylation data from four studies in a unified cohort, totaling 727 ccRCC tumors, to compare published methylation-based classification methods and to investigate their relevance for treatment response. The four classifications methods we evaluated were all associated with overall survival in the unified cohort, and their discriminative performance was comparable. The Sato et al. method^14^ identified the smallest hyper-methylated group, which carried the highest risk of death, and its prognostic value was confined to stage III-IV tumors, independently of stage in multivariable analysis. This suggests that methylation subtyping is most informative in patients at high risk of progression, precisely those who are candidates for systemic therapy. The lower proportion of women among hyper-methylated tumors is an intriguing observation that warrants further investigation.

Hyper-methylated tumors were characterized by a larger fraction of cycling tumor cells (PCNA and MKI67-positive) and by the activation of proliferation programs, including the MYC, E2F and G2/M checkpoint gene sets ^42,62^. Two independent mTOR perturbation signatures, together with the mTORC1 hallmark gene set, consistently indicated higher mTOR activity in these tumors. The unfolded protein response was also activated, a process that has been linked to tumor progression and treatment resistance ^63,64^. Conversely, hypo-methylated tumors contained higher fractions of endothelial tip cells, in agreement with the recent work of Lu et al. ^37^. Hyper-methylated tumors also showed reduced activity of homeobox TFs, including NANOG, PBX3 and POU5F1, and of GATA factors whose loss has been associated with aggressiveness in ccRCC ^47–49^. This observation is consistent with the over-representation of homeobox genes among the loci targeted by the Sato et al. signature probes ^14^, and with the association between low homeobox gene expression and poor prognosis in ccRCC ^65^. Finally, the over-expression of the de novo methyltransferase DNMT3B and the under-expression of the demethylase TET2 suggest a mechanism for the hyper-methylated phenotype ^66,67^. This hypothesis remains to be tested functionally, for instance by modulating these enzymes in ccRCC models.

Because methylation profiling is rarely performed in clinical trials, we developed a classifier that infers methylation subtypes from gene expression data. Its gene-ratio design makes it independent of normalization and applicable across platforms, which allowed us to apply it without further processing to four independent cohorts. The classifier was optimized for specificity: predicted hyper-methylated tumors were highly reliable, whereas about one third of hyper-methylated tumors were classified as hypo-methylated. This misclassification is expected to attenuate rather than inflate differences between predicted subtypes, suggesting that the associations observed in treated cohorts may underestimate the true effect of methylation status.

The association between predicted hyper-methylation and poor response to sunitinib was consistent across primary tumors from three cohorts and metastases from IMmotion151. A plausible explanation is that hyper-methylated tumors contain fewer endothelial tip cells, the main drivers of sprouting angiogenesis and presumed targets of VEGF receptor inhibition. This interpretation agrees with the IMmotion151 molecular subsets, in which angiogenic tumors responded better to sunitinib, whereas proliferative tumors had poor outcomes but benefited from atezolizumab plus bevacizumab ^68^. Our results extend these observations by linking this angiogenic-proliferative axis to a DNA methylation phenotype. They also echo earlier work reporting a CpG island methylator phenotype among the ccrcc1 and ccrcc4 subgroups, which respond poorly to sunitinib ^38^. Unlike the classifiers of Lu et al. ^37^, our subtypes predicted from gene expression retained their association with PFS in independent cohorts. Notably, the immune composition and TLS scores of the two subtypes were similar, which may explain why methylation subtypes were not associated with response to immune checkpoint inhibition alone.

Within the randomized IMmotion151 and IMmotion150 trials, atezolizumab plus bevacizumab was associated with longer PFS than sunitinib in patients with hyper-methylated primary tumors. In contrast, sunitinib was more effective in patients with hypo-methylated metastases. These findings raise the possibility that methylation subtypes could help choose between anti-angiogenic monotherapy and immune checkpoint-based combinations. The prospective BIONIKK trial has shown that such molecularly guided treatment selection is feasible in metastatic ccRCC ^29^.

Several limitations should be considered. First, all analyses were retrospective, and treatment comparisons within predicted subtypes are subgroup analyses of randomized trials, which require confirmation in independent cohorts. Second, results in metastatic samples relied on a single cohort. Third, sunitinib was the only TKI evaluated, and current first-line regimens combine immune checkpoint inhibitors with TKIs or with other immunotherapies ^5,6^. The relevance of methylation subtypes for these combinations therefore remains to be established. Despite these limitations, our study shows that a DNA methylation phenotype, inferable from routine transcriptomic data, captures proliferative and angiogenic features of ccRCC that are associated with survival and treatment response. If validated prospectively, this approach could contribute to the personalization of systemic therapy in metastatic ccRCC.

## Methods

### Ethics

This study used only publicly available or controlled-access datasets, which had been collected with patient consent and ethics approval as described in the original publications. Access to controlled data from the European Genome-phenome Archive (EGA) was granted by the respective data access committees. No additional ethics approval was required.

### DNA methylation data

Methyl-array data from four independent ccRCC cohorts were collected (Fig. S1), comprising data from The Cancer Genome Atlas (TCGA) (450k Illumina array, N = 318)^13^, Clinical Proteomic Tumor Analysis Consortium (CPTAC) (methyl-array EPIC Illumina array, N = 196)^15,16^, Sato et al. (450k Illumina array, N = 111)^14^ and Evelönn et al. (450k Illumina array, N = 132)^32^. For TCGA patients profiled on more than one platform, a single sample was retained, with priority to the 450k Illumina array to maximize probe overlap across cohorts.

Probes with more than 15% missing values were removed, followed by samples with more than 20% missing values. Remaining missing values were imputed by the knn() function of the VIM R package (v. 6.2.2). Batch effects between cohorts were corrected with the ComBat algorithm of the R/Bioconductor package sva (v. 3.42.0), using the cohort as the batch variable. The final unified dataset comprised 727 tumors.

### Gene expression data

Gene expression data from seven cohorts, totaling 2,072 samples, were used in our study: The Cancer Genome Atlas (TCGA) (N = 301, RNA-Seq, in transcripts per million) ^13^, Clinical Proteomic Tumor Analysis Consortium (CPTAC) (N = 187, RNA-Seq, in transcripts per million)^15,16^, the Sato et al. dataset (N = 101, micro-array)^14^, the Braun et al. dataset (N = 309, RNA-Seq, normalized transcripts per million)^11^, the Beuselinck et al. dataset (N = 53, micro-array)^38^, the Motzer et al. dataset (N = 823)^10^ and the Banchereau et al. dataset (N = 247)^59^.

The TCGA, CPTAC and Sato et al. cohorts, with matched methylation data (N = 589), were used for biological characterization and classifier development. The four other cohorts, comprising patients with advanced ccRCC treated with systemic therapies, were used to evaluate associations with treatment response.

The Braun et al. dataset includes primary tumors and metastases from patients enrolled in CheckMate 009 (NCT01358721), CheckMate 010 (NCT01354431) and CheckMate 025 (NCT01668784) trials. These patients had progressed after one to three prior therapies, including at least one anti-angiogenic agent, and received nivolumab (anti-PD-1) or everolimus (mTOR inhibitor) (EGAC00001001519, EGAC00001001520, EGAC00001001521). Only nivolumab-treated patients were analyzed (n = 180). The Beuselinck et al. dataset comprises primary tumors from patients with metastatic ccRCC treated with first-line sunitinib (N = 53). The Motzer et al. dataset comprises primary tumors and metastases from the phase 3 IMmotion151 trial (NCT02420821) (N = 823). Patients were randomized to sunitinib or atezolizumab (anti-PD-L1) plus bevacizumab (anti-VEGF) (EGAC00001001813). The Banchereau et al. dataset consists in samples from the phase 2 IMmotion150 trial (NCT01984242) (N = 247). Patients were randomized to sunitinib, atezolizumab plus bevacizumab or atezolizumab alone (EGAC00001001748).

### Clustering of tumors into DNA methylation subtypes

Four published methods were applied to beta values of the unified methylation dataset. The Arai et al. method divided ccRCC tumors into a CpG island methylator phenotype-positive group (CIMP+, hyper-methylated, “High”) and a CIMP-group (hypo-methylated, “Low”). The method uses hierarchical clustering (Ward’s method, Euclidean distance) of 801 probes originally selected on the 27K Illumina array.

The Sato et al. method split ccRCC tumors by hierarchical clustering (agglomeration method Ward.D2 agglomerative method, Euclidean distance) based on 1,672 probes selected on the 450K Illumina array. Tumors were divided into two groups, and a three-cluster split was evaluated separately. The Evelönn et al. method divided ccRCC tumors using a classification score based on 172 probes. The Lu et al., method classified ccRCC tumors into two subtypes based on 58 probes, using the iMES R package (https://github.com/xlucpu/iMES) ^37^. Because the method does not accept missing values, five probes with missing values were imputed by k-nearest neighbors (k = 10).

After cross-platform harmonization and missing values filtering, 531 of 801 probes (Arai et al), 1,006 out 1,672 probes (Sato et al.) and 148 of 172 probes (Evelönn et al.). Classifications obtained with these reduced probe sets were concordant with those obtained with the complete signatures.

### Differential expression, cell type deconvolution and pathway analyses

Gene expression was compared between methylation subtypes with two-sided Wilcoxon rank-sum tests in the TCGA and Sato et al. datasets, for the 20,085 genes shared by both datasets.

Cell type fractions were estimated with CIBERSORTx (https://cibersortx.stanford.edu, v1.0, 12/21/2019)^69^ using as reference a refined and relabeled ccRCC single-cell RNA-seq dataset^42,70^.

Single-sample Gene Set Enrichment Analysis (ssGSEA) of the MSigDB hallmark (H) and oncogenic signatures (C6) collections (Human MSigDB v2023.1.Hs)^43,44^ was performed with the GSVA R/Bioconductor package (v. 1.42.0).

Transcription factor (TFs) activities were inferred with the VIPER algorithm^45^ (msviper() function of the viper R/Bioconductor package v. 1.28.0; minsize = 15, adaptive.size = F, pleiotropy = TRUE). The analysis used 271 TF regulons of confidence levels A-C from the DoRothEA R/Bioconductor package (v. 1.12.0)^71^, restricted to TFs with at least 15 measured transcripts. The input signature was the log2 fold change in gene expression between hyper-methylated and hypo-methylated tumors.

A tertiary lymphoid structure (TLS) score was computed from a seven-gene signature (CCL19, CCL21, CXCL13, CCR7, CXCR5, SELL, LAMP3) as previously described^39^. Raw scores were compared between subtypes with two-sided Wilcoxon rank-sum tests. For visualization only, scores were dichotomized at the median.

Cell type deconvolution, ssGSEA and TF activity inference analyzes were performed separately in each of the three cohorts. Cell fractions and ssGSEA values were standardized (z-scores) within each cohort. P values were combined across cohorts with Stouffer’s method, and NES values were averaged for visualization. Only features significant in all three cohorts were retained.

### Gene expression-based classifier

A classifier predicting methylation subtypes from gene expression data was developed with the Cross-Platform Omics Prediction (CPOP) algorithm^58^. CPOP builds penalized regression models on pairwise log-ratios of gene expression and selects ratios with stable effects across training datasets. The model was trained on the TCGA (RNA-seq) and Sato et al. (microarray) datasets, chosen to maximize sample size and to combine distinct profiling platforms. It was evaluated on the independent CPTAC dataset, which was not used for model training or hyperparameter selection.

Feature selection proceeded in two steps, both performed on the training datasets only. Firstly, differentially expressed genes (DEGs) between methylation subtypes were identified in each training dataset (two-sided Wilcoxon rank-sum tests), and only DEGs shared by both datasets were retained. Secondly, the number of DEGs (100, 125, 150, 175 or 200) and the number of features provided to CPOP (10, 20, 30 or 40) were optimized by grid search to minimize the false-positive rate, defined as hypo-methylated tumors misclassified as hyper-methylated.

The final model, built on 40 features selected from the top 125 shared DEGs, yielded a nine-gene ratio signature after removal of correlated features (CDCA3/IGF2BP3, MYBL2/PCCA, MCF2L/PDE2A, MCF2L/SRPX2, CDCA3/MCF2L, RNF152/UBE2C, MCF2L/PTPRH, and MCF2L/TNNT1). It consists of two sub-models, one trained on each training dataset. For a new sample, the probabilities from the two sub-models are averaged. Samples with an averaged probability above 0.5 are classified as hypo-methylated, and the others as hyper-methylated. No normalization was applied beyond that of the original publications, as gene ratios are insensitive to sample-level scaling.

The final model, which was targeting the selection of 40 features from 125 DEGs, yielded a 9-gene-ratio signature after internal gene ratio calculation from gene expression values and correlated features removal. The ratios retained were AURKB/MCF2L, CDCA3/IGF2BP3, MYBL2/PCCA, MCF2L/PDE2A, MCF2L/SRPX2, CDCA3/MCF2L, RNF152/UBE2C, MCF2L/PTPRH, and MCF2L/TNNT1. At the end, the classification model consisted of two internal sub-models, trained on TCGA and Sato et al datasets. To classify a new sample, each sub-model generated a probability score, and the final classification was determined by averaging these probabilities. Samples with an averaged probability greater than 0.5 were designated as hypo-methylated, whereas those with a probability of 0.5 or lower were designated as hyper-methylated. No additional normalization procedures (e.g., min-max scaling or Z-score transformation) were applied beyond those used in the original publications. This approach demonstrated strong robustness in handling heterogeneity from gene expression data.

Performance in CPTAC was assessed with the hyper-methylated subtype as the positive class. The AUC-ROC was computed from averaged probabilities. Sensitivity, specificity, positive and negative predictive values and accuracy are reported with Wilson 95% confidence intervals (CIs). The Matthews correlation coefficient (MCC) is reported with a 95% CI from 2,000 bootstrap resamples.

### Survival analysis

Survival curves were estimated with the Kaplan-Meier method and compared with two-sided log-rank tests, using the survival (v. 3.4-0) and survminer (v. 0.4.9) R packages. Hazard ratios (HRs) and 95% CIs were estimated with Cox proportional hazards models (Efron method for ties), with the hypo-methylated subtype as reference. The independent association of methylation subtype and tumor stage with OS was assessed with a multivariable Cox model. The discriminative ability of each methylation classification was quantified with Harrell’s C-index, with 95% CIs from 1,000 bootstrap resamples.

### Statistical analysis

Continuous variables were compared between two groups with two-sided Wilcoxon rank-sum tests. P values were adjusted for multiple testing with the Benjamini-Hochberg method where indicated. Categorical variables were compared with Pearson’s chi-squared test, or with Fisher’s exact test when any expected count was below 5. All analyses were performed in R (v. 4.1.2) with Bioconductor/BiocManager (v. 3.14).

## List of abbreviations

AUC-ROC: area under the curve of the receiver operating characteristic curve
ccRCC: clear cell renal cell carcinoma
CB: clinical benefitl
CI: confidence interval
CIMP: CpG island methylator phenotype
CPOP: Cross-Platform Omics Prediction
CpG: 5′-cytosine-phosphate-guanine-3’
CPTAC: Clinical Proteomic Tumor Analysis Consortium
CR: complete response
DEG: differentially expressed genes
DNMT: DNA methyltransferase
EGA: European Genome-phenome Archive
FDA: U.S. Food and Drug Administration
HR: hazard ratio
ICB: intermediate clinical benefit
ICI: immune checkpoint inhibitor
IHC: immunohistochemistry
ITH: intratumoral heterogeneity
MCC: Matthews correlation coefficient
MHC: major histocompatibility complex
MSigDB: Molecular Signatures Database
MSKCC: Memorial Sloan Kettering Cancer Center
mTOR: mammalian target of rapamycin
NAT: normal adjacent tissue
NCB: non-clinical benefit
NE: inevaluable
NES: normalized enrichment score
NPV: negative predictive value
NSCLC: non-small cell lung cancer
OS: overall survival
PD: progressive disease
PD-1: programmed cell death protein 1
PD-L1: programmed death-ligand 1
PFS: progression-free survival
PPV: positive predictive value
PR: partial response
pRCC: papillary renal cell carcinoma
RCC: renal cell carcinoma
RECIST: Response Evaluation Criteria in Solid Tumors
SD: stable disease
ssGSEA: single-sample gene set enrichment analysis
TCGA: The Cancer Genome Atlas
TET: Ten-Eleven Translocation (DNA demethylases)
TF: transcription factor
TKI: tyrosine kinase inhibitors
TLS: tertiary lymphoid structure
TMB: tumor mutation burden
TME: tumor microenvironment
TPM: transcripts per million
VEGF: vascular endothelial growth factor
VIPER: Virtual Inference of Protein-activity by Enriched Regulon analysis

## List of Supplementary Materials

Fig S1 to S7

Supplementary Tables.

## Acknowledgments

Most of the computations presented in this paper were performed using the GRICAD infrastructure (https://gricad.univ-grenoble-alpes.fr), which is supported by Grenoble research communities. We acknowledge Genentech for the access to Motzer et al. (EGAC00001001813) and Banchereau et al. (EGAC00001001748) datasets of normalized gene expression data and associated clinical features.

## Author contributions

F.J., C.B. and D.P. conceived and designed the study. F.J. developed the methodology, collected and curated the data, performed the computational analyses and prepared the figures, with contributions from S.S., H.A. and P.B. to the investigation. S.N.S. and A.L. provided clinical expertise and contributed to the interpretation of the treatment response analyses. C.B. and D.P. supervised the project and acquired funding. F.J., C.B. and D.P. wrote the manuscript. All authors critically reviewed and edited the manuscript and approved the final version.

## Data and materials availability

The TCGAbiolinks R package (v. 2.22.4) was used to download TCGA and CPTAC datasets (methyl-array and RNA-Seq). The Sato et al. datasets were collected from ArrayExpress (E-MTAB-2007). The Beuselinck et al. datasets were collected from ArrayExpress (E-MTAB-3267). The Evelönn et al. and the Braun et al. datasets were collected from the data sources of the articles. The Motzer et al. normalized gene expression data (TPM) generated by RNA-Seq and clinical data of 823 participants were obtained from the European Genome-phenome Archive (EGA) (EGAC00001001813). The Banchereau et al. normalized gene expression data (TPM) generated by RNA-Seq and clinical data were obtained from the EGA (EGAC00001001748). All results are available in the main text and in the supplementary materials. A Zenodo repository provides data and code, implemented in R language with Jupyter Notebooks and embedded in a Singularity container, to reproduce the entire analysis workflow (https://doi.org/10.5281/zenodo.15052573).

